# Revisiting and expanding the meta-analysis of variation: The log coefficient of variation ratio, lnCVR

**DOI:** 10.1101/2020.01.06.896522

**Authors:** Alistair M. Senior, Wolfgang Viechtbauer, Shinichi Nakagawa

## Abstract

Meta-analyses are frequently used to quantify the difference in the average values of two groups (e.g., control and experimental treatment groups), but examine the difference in the variability (variance) of two groups. For such comparisons, the two relatively new effect size statistics, namely the log-transformed ‘variability ratio’ (the ratio of two standard deviations; lnVR) and the log-transformed ‘CV ratio’ (the ratio of two coefficients of variation; lnCVR) are useful. In practice, lnCVR may be of most use because a treatment may affect the mean and the variance simultaneously. We review current, and propose new, estimators for lnCVR and lnVR. We also present methods for use when the two groups are dependent (e.g., for cross-over and pre-test-post-test designs). A simulation study evaluated the performance of these estimators and we make recommendations about which estimators one should use to minimise bias. We also present two worked examples that illustrate the importance of accounting for the dependence of the two groups. We found that the degree to which dependence is accounted for in the sampling variance estimates can impact heterogeneity parameters such as *τ*^2^ (i.e., the between-study variance) and *I*^2^ (i.e., the proportion of the total variability due to between-study variance), and even the overall effect, and in turn qualitative interpretations. Meta-analytic comparison of the variability between two groups enables us to ask completely new questions and to gain fresh insights from existing datasets. We encourage researchers to take advantage of these convenient new effect size measures for the meta-analysis of variation.

## 1. INTRODUCTION

Meta-analysis is often used to evaluate studies comparing the average of two groups. These are usually treatment groups in an experiment/trial, one being a concurrent control, but may also represent naturally occurring groups (e.g., different sexes). The standardised mean difference (SMD; also known as Cohen’s *d* and its associated derivatives), which is the difference between group means divided by the within-study variability, is a commonly-used effect size measure for this purpose^1^. SMD is popular because it is ‘unitless’, meaning it can be used to compare the results of studies that report outcomes in different units^2^ A similar unitless effect size measure, which can also be used to compare the means of two groups, is the logarithm of the ratio of the means of the groups. This effect size measure is known as the ratio of means in medicine (ROM^3^) and the log response ratio in ecology and evolution (lnRR^4^). Throughout we follow the lnRR notation as this will help to draw parallels with other effect size measures as we progress, but the reader should not be confused with the (logarithm of) risk ratio, which is also sometimes denoted (ln)RR. Surveys have shown that lnRR is the most widely used effect size measure in ecology and evolution^5–7^ Moreover, SMD and lnRR collectively account for over half of all meta-analyses in ecology^6,7^ meaning comparisons between group means is the most widespread aim of meta-analysis in this field. SMD also seems to be among the most used standardised effect statistics in the medical and social sciences^8^.

Two groups may not only differ in terms of their means, but also their variances^9^. At the most basic level, experimental treatments may directly increase or decrease the total amount of variance in a system due to inter-individual variability in response. Many biological systems also appear to display a mean-variance relationship^10–12^; most commonly, increasing averages are associated with increasing variances. Perhaps the most well-known example of a biological mean-variance relationship comes from ecology and is known as Taylor’s Law. This ‘law’ has been widely observed, and states that as mean population density increases, variance in population density also increases^13,14^. Where mean-variance relationships are present, a treatment may indirectly cause groups to have differing variances by altering the mean.

Nakagawa, Poulin, Mengersen, et al.^15^ proposed a number of methods that allow the user to test for differences in the variance of groups meta-analytically. Among the methods proposed, the logarithm of the ratio of the standard deviations (SDs), named log ‘variability ratio’ (lnVR) and the logarithm of the ratio of the coefficients of variation (CV), termed the log ‘CV ratio’ (lnCVR), are most readily integrated into the standard meta-analytic paradigm; i.e. a contrast-based model using an effect size that corresponds to an effect relative to a concurrent control^16,17^. Of the two, lnCVR is perhaps the more useful measure where a mean-variance relationship is likely to exist. Nakagawa, Poulin, Mengersen, et al.^15^ highlight that meta-analysing variation may be used to answer completely novel questions, but it can also be used to provide fresh insights into the topics on which a meta-analysis of means was already conducted. Indeed, lnCVR has already been applied in such diverse fields as ecology^18^, evolution^19^, agriculture^20^, health^17^, and the social sciences^21^. It is important to note that lnCVR (and also lnVR) require the same data to calculate as is already needed for computing SMD or lnRR values.

Our aims in this paper are threefold. First, we review existing and propose new estimators for lnCVR and its sampling error variance. These include, for the first time, estimators of the sampling variance when the two groups (treatment and control) are not independent (as may occur, for example, in cross-over trials or in paired, single-subject, or pre-test-post-test designs). Second, we conduct a simulation study to investigate the performance of the different estimators. Finally, we present two case studies using these techniques, and illustrate the importance of accounting for dependence between the two treatment groups in the estimation of sampling variation and other heterogeneity parameters (e.g., *τ*^2^, the between-study variance, and *I*^2^^22^).

## 2. METHODS

### 2.1 Point estimators when groups are independent

Let *x_T_* ~ *N*(*μ_T_, σ_T_*) and *x_c_* ~ *N*(*μ_C_, σ_c_*) denote normally distributed random variables with true means (i.e., expected values) given by *μ_T_* and *μ_C_* and true standard deviations *σ_T_* and *σ_C_*. For independent random samples based on these variables (e.g., representing some outcome of interest measured in a treatment and control group) of size *n_T_* and *n_C_*, let 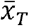 and 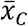 denote the respective sample means and *s_T_* and *s_C_* the corresponding standard deviations for the two groups. Then comparisons between the means, variances and coefficients of variation for two groups can be made using the lnRR, lnVR and lnCVR effect size measures, respectively. “Naïve” estimators of these effect statistics are:

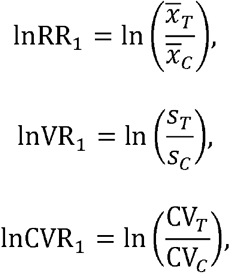

where ln denotes the natural logarithm and 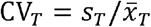 and 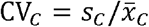 denote the coefficients of variation in the treatment and control group, respectively.

While these naïve estimators are consistent and asymptotically unbiased, we can add corrections for the sample size based on a second-order Taylor expansion (also known as, the second order delta method) for each statistic^15,23,24^. For the lnRR, Lajeunesse^23^ demonstrated such a correction is important to obtain unbiased estimation especially when sample size is small;

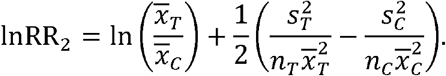

Similarly, for the lnVR, Nakagawa, Poulin, Mengersen, et al.^15^ proposed:

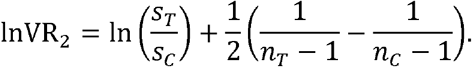

Combing lnRR2 and lnVR2, one obtains:

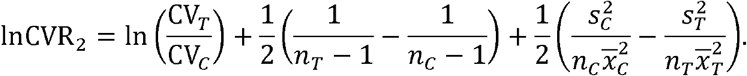

Note that Nakagawa, Poulin, Mengersen, et al.^15^ originally suggested the an estimator of lnCVR that missed the bias correction pertaining to lnRR (i.e. 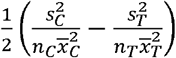). We also note that an alternative estimator of lnCVR could also be obtained based on 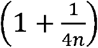 CV, which it has been suggested acts as a ‘rough’ bias correction for the CV (e.g.^25^). However, this estimator is not recommended here, and it does not perform well (see Supplementary Materials S1, Text S1).

### 2.2 Dispersion estimators when the two groups are independent

The original estimators of the sampling (error) variance for lnRR^4^ and lnVR^15^ are based on the first-order Taylor expansion; they are respectively:

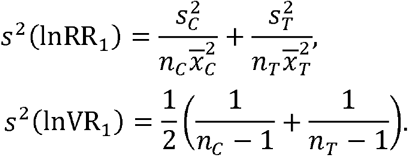

Based on these, for lnCVR Nakagawa, Poulin, Mengersen, et al.^15^ proposed:

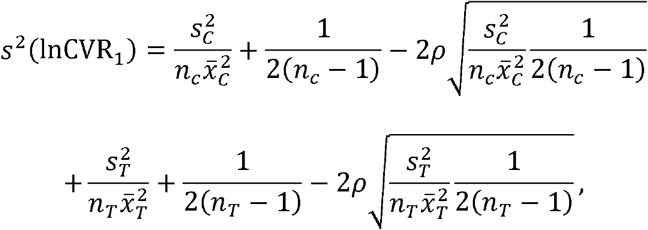

where *ρ* is the correlation between the log mean and log SD. Nakagawa, Poulin, Mengersen, et al.^15^ suggested that *ρ* can be estimated based on the correlation between the log sample mean and log sample SD across the studies included in a meta-analysis. However, in doing so one risks conflating within- and between-study correlation (i.e., the correlation in the bivariate sampling distribution of the sample mean and sample SD could be very different than the correlation of the true means and SDs across studies). In fact, for observations that come from an underlying population distribution that is symmetric (e.g. a normal distribution), the sample mean and variance are uncorrelated^26^. Thus, for the case considered here where *ρ* = 0 the equation above simplifies to:

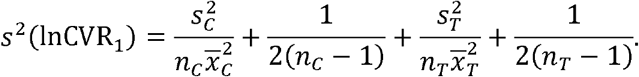

As a better estimator for the sampling variance of lnRR, Lajeunesse^23^ derived and tested the following sampling variance based on the second-order Taylor expansion:

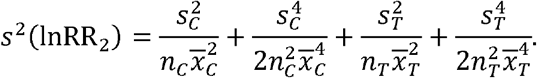

Similarly, we can derive the following sampling variance for lnVR based on the second-order Taylor expansion as:

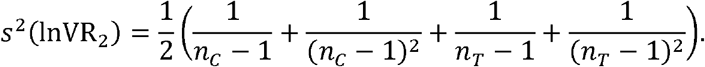

Accordingly, the complete estimator of the sampling variance for lnCVR, based on *s*^2^(lnRR_2_) and *s*^2^(lnVR_2_) is:

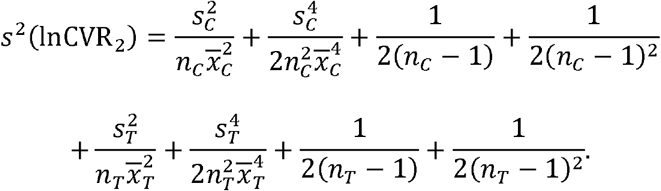

In the supplementary materials, we propose estimators of the sampling covariance based on the above, which can be used when multiple treatment groups are contrasted with the same control^27^ (see Supplementary Materials S1, Text S2)

### 2.3 Point estimators when groups are dependent

Due to experimental design, control and treatment groups are often not independent of one another. A clear example of this dependency is in the case of a cross-over design where the same individuals are subjected to both control and experimental treatments at two different time points. The point estimates given above will perform the same way regardless of whether we are dealing with independent or dependent groups. In cross-over studies, however, *n_T_* = *n_c_* ≡ *n*, unless dropouts are included in a pre-post design, in which case we recommend that *n* = *n*_post_ (i.e. the sample size in the post-treatment condition) is used. This is because the correlation between pre and post-treatment measurements can only be calculated based on *n*, which assumes *n_T_* = *n_c_* (see the next section). We can rewrite the dependent cases of lnRR_1_ and lnRR_2_ as:

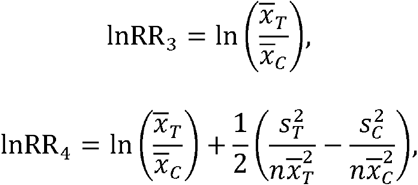

where subscripts 3 and 4 indicate the naïve estimator and estimator based the second-order Taylor expansion, respectively. Similarly, for lnVR and lnCVR, we have:

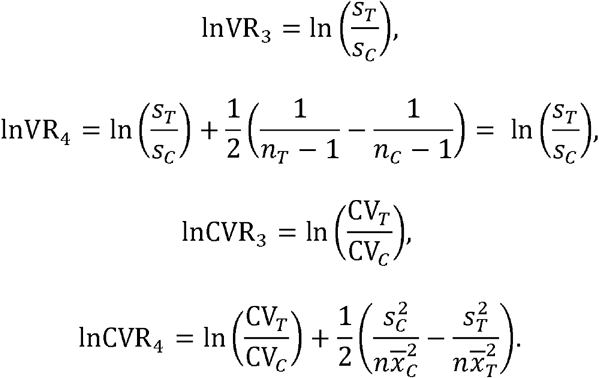

### 2.4 Dispersion estimators when the two groups are dependent

In dependent cases estimates of the sampling variance need to account for the correlation between measurements from the same replicates on the two occasions (i.e. cross-correlation^28^). Based on the first-order Taylor expansion, the sampling variance for lnRR is:

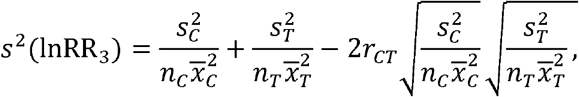

where *r_CT_* is a cross-context correlation value estimated from the two sets of measurements on the same replicate when they are under the control and treatment conditions^29^ As discussed above for dependent studies *n_τ_ = n_c_ ≡ n* meaning *s*^2^(lnRR_3_) simplifies to:

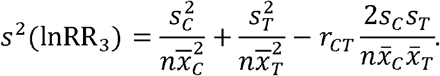

If based on the second-order Taylor expansion^23^, the estimator of the sampling variance for lnRR is:

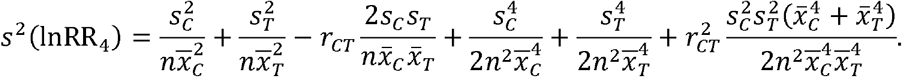

We can also derive the sampling variance for dependent cases of lnVR based on the first- order Taylor expansion as:

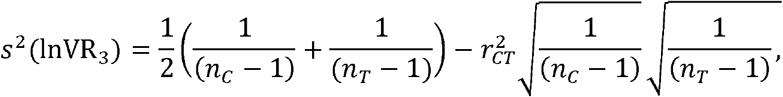

which, where *n_T_* = *n_c_ = n*, simplifies to:

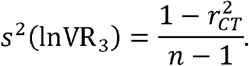

Based on the second-order Taylor expansion, we have the sampling variance for dependent cases of lnVR as:

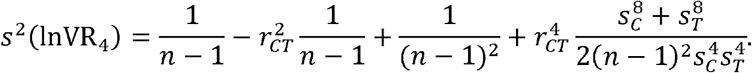

From the sampling variances for lnRR and lnVR, we have the sampling variance for lnCVR with first- and second-order Taylor expansion as:

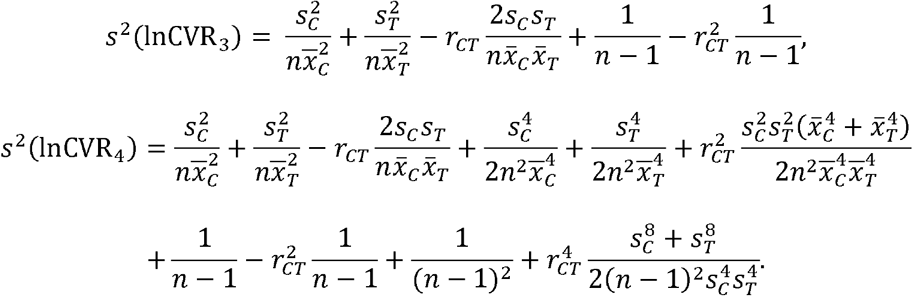

Note that, where *r* is positive the estimated sample variance for a dependent estimator will be smaller than its independent equivalent, but that as *r* shrinks to 0 the dependent case converges on the independent; e.g. assuming *n_c_* = *n_T_*, where *r* > 0, *s*^2^(lnCVR_3_) < s^2^(lnCVR_1_), but where *r* = 0, *s*^2^(lnCVR_3_) = *s*^2^(lnCVR_1_).

## 3. SIMULATION

### 3.1 Simulation study design

We simulated a two-group experiment/trial, where a pair of groups is based on *n_T_* and *n_C_* random samples drawn from populations under an experimental treatment and control conditions. The treatment and control populations have means *μ_T_* and *μ_C_* and standard deviations (SDs) *σ_T_* and *σ_C_*, respectively. The *i*th sample in each group, *y_Ti_* (*i* = 1 … *n_T_*) and *y_Ci_* (*i* = 1 … *n_C_*) was drawn from a bivariate normal distribution as follows:

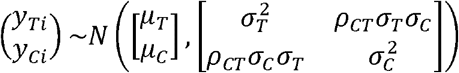

Where 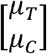 are the population means of the two groups, 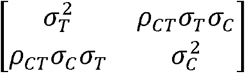 is a variance co-variance matrix specifying the variances of the two groups with *p_CT_* giving the degree of correlation among the *i*th samples in the two groups and all other parameters are as above. When where *p_CT_* ≠ 0 the *i*th data in the two groups are correlated (i.e. dependent or paired samples as in a cross-over design).

In all simulations, *μ_C_* = 100 and *σ_C_* = 20, which across the parameters tested ensures positive sample means (required for log transformation). We explored values of *μ_T_* ranging between *μ_C_* × *e*^-0.5^ and *μ_C_* × *e*^0.5^ and values of *σ_T_* ranging between *σ_C_* × *e*^0.5^ and *σ_C_* × *e*^0.5^, meaning the ln(*μ_T_*/ *μ_C_*) and ln(*σ_T_*/ *σ_C_*) is between −0.5 and 0.5. All combinations were explored and where ln(*μ_T_* / *μ_C_*) = ln(*σ_T_* / *σ_C_*) the coefficient of variance (CV) of the two groups will be identical. We explored *n_C_* = 8, 16 and 42, with *n_C_* = *n_T_* and, with *n_C_* < *n_T_* (independent case). We also explored *p_CT_* = 0 and *p_CT_* = 0.8. For each set of parameters, we simulated 100,000 experiments.

Based on the sample means and SDs of each simulated experiment, we calculated lnCVR_1_ and lnCVR_2_ for independent cases (*ρ_CT_* ≠ 0) and lnCVR_3_ and lnCVR_4_ for dependent cases (*ρ_CT_* ≠ 0). We also calculated the sampling variance estimators *s*^2^(lnCVR_1_) and *s*^2^(lnCVR_2_) where *ρ_CT_* ≠ 0, and *s*^2^(lnCVR_3_) and *s*^2^(lnCVR_4_) where *ρ_CT_* ≠ 0. We calculated bias in the *i*th estimator as:

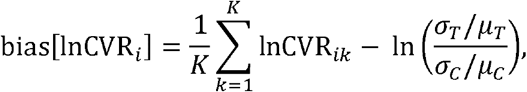

where *k* is the *k*th value of *K* (here 100,000) simulated values of lnCVR*_i_* (*k* = 1… *K*). This bias can be interpreted as the mean deviation of the *i*th estimator of lnCVR from the true population value. We calculated bias in sampling variance estimator *i* as:

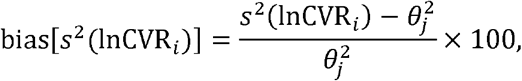

where *s*^2^(lnCVR*_i_*) is the value of the *i*th sampling variance based on the simulated population statistics and sample sizes and *θ*_*j*_ is the SD among *K* simulated effect sizes estimated using estimator *j*. This bias can be interpreted as the percentage by which the sampling variance estimator deviates from the true value (i.e. 100 = the estimator is twice the true value). We calculated coverage as the proportion of 95% confidence intervals (CIs) that include In 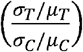. For a combination of the *j*th effect size estimator (lnCVR*_j_*) and *i*th sampling variance *s*^2^(lnCVR*_i_*), 95% CIs were constructed as:

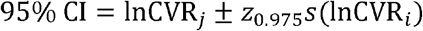

where lnCVR*j* is the estimated effect size for the simulated sample, *s*(lnCVR*_i_*) an estimate of the standard error (SE; the square root of the estimated sampling variance), and z_0.975_ is the function of the 0.975^th^ quantile of a *z* distribution (approx. 1.96). Simulations and analyses were performed in R v3.5.1;, and using the ‘mvrnorm’ function in the *MASS* package^31^. All data and code presented in this manuscript can be found at (https://github.com/AlistairMcNairSenior/lnCVR_Estimators_Sim).

## 3.2 Simulation results

We begin with the case where the two groups are independent (*ρ_CT_* = 0). Figure 1 shows bias in the estimated effect as a function of sample size and the log the ratio of the means and SDs in the two groups. Across the diagonal elements of each plot (black-dashed line) the underlying CV of the two populations is identical (even if the means and SDs differ; 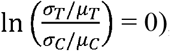), elements above the line correspond to the CV of the treatment population being greater than that of the control group 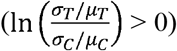, and elements below the line the opposite 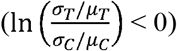. lnCVR_1_ overestimates positive effects and slightly under-estimate negative effects, with bias being most profound where the sample size is small. lnCVR_2_, on the other hand, displays no systematic bias. Figure 2 shows the results where the sample size of the treatment group is ~25% greater than that of the control group. lnCVR_1_ showed severe upward bias, especially where the sample size was small, where as lnCVR_2_ performed with only very minor upward bias, which all but disappeared for larger sample sizes. Given that lnCVR_2_ was determined to be the most accurate estimator of the effect, we proceeded to explore how lnCVR_2_ performed in conjunction with different estimators of sampling variance.

**Figure 1:**
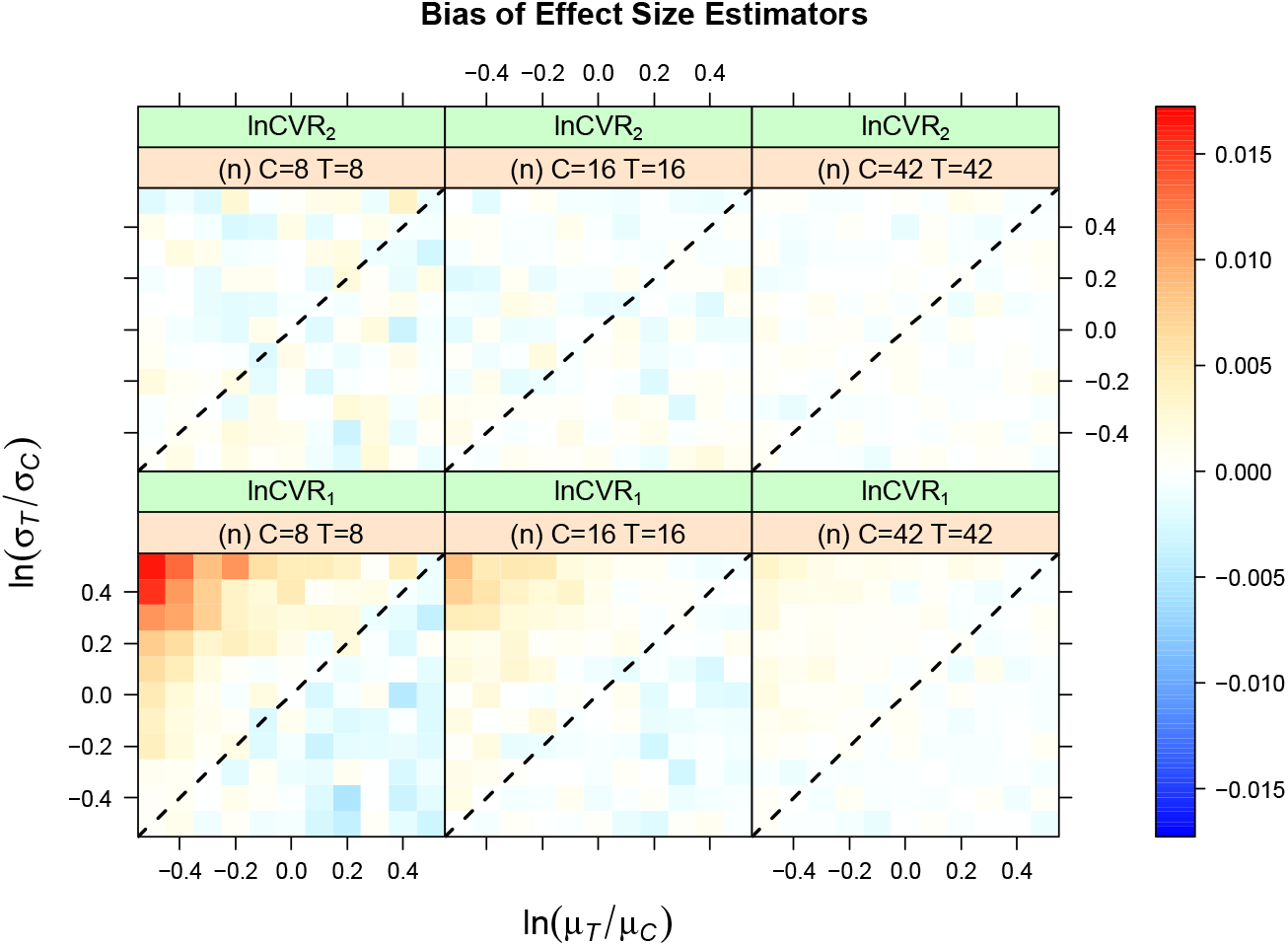
Bias in effect size estimators of lnCVR as a function of the log ratio of population means (*x*-axis), SDs (*y*-axis) and sample size (balanced) for the case of independent treatment and control group data (*ρ_CT_* = 0). Black dashed line indicates no effect (i.e., lnCVR = 0).

The first sampling variance estimator *s*^2^(lnCVR_1_) underestimated the variance among simulated values of lnCVR_2_, particularly where the sample size was small (Figure 3). Biases for *s*^2^(lnCVR_2_) were minimal, although there was some very slight upward bias for small sample sizes and large positive effects (Figure 3). The coverage of 95% CIs for *s*^2^(lnCVR_1_) and *s*^2^(lnCVR_2_) (paired with lnCVR_2_) are shown in Figure 4. *s*^2^(lnCVR_1_) generated CIs that were too narrow at smaller sample sizes, whereas again *s*^2^(lnCVR_2_) performed with little bias. At larger sample sizes coverage was much closer to the nominal level (Figure 4), although *s*^2^(lnCVR_2_) still performed more accurately. The same patterns of performance were observed for the case where *n_C_* < *n_T_* (Supplementary Figures S1 and S2).

**Figure 2:**
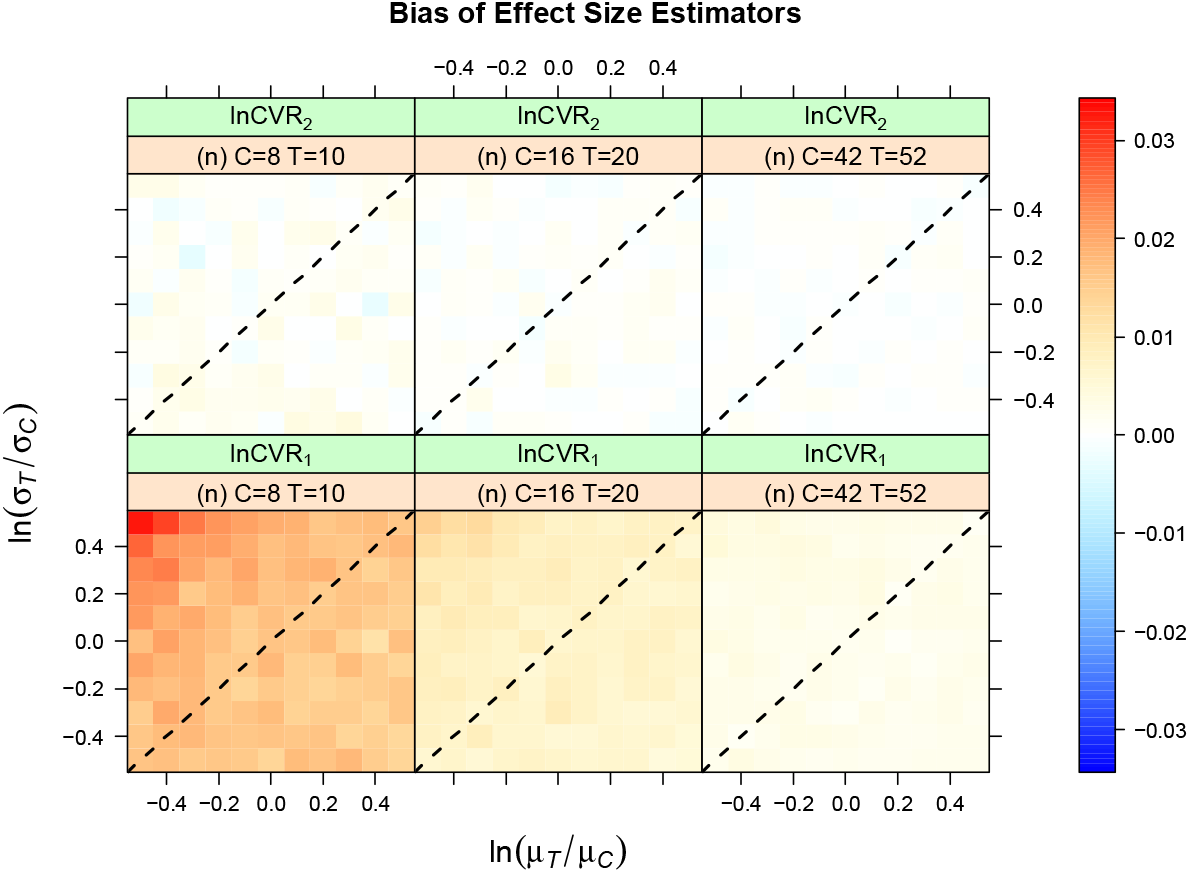
Bias in effect size estimators of lnCVR as a function of the log ratio of population means (*x*-axis), SDs (*y*-axis) and sample size (unbalanced) for the case of independent treatment and control group data (*ρ_CT_* = 0). Black dashed line indicates no effect (i.e., lnCVR = 0).

**Figure 3:**
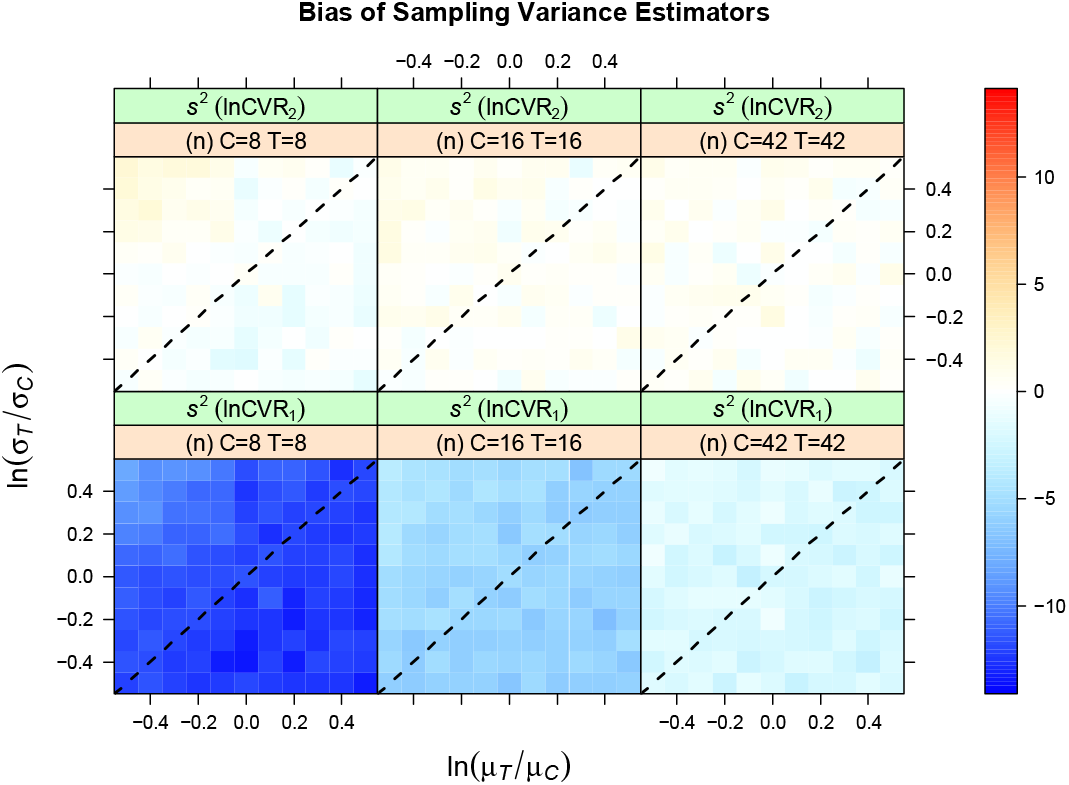
Bias in sampling variance estimators of lnCVR as a function of the log ratio of population means (*x*-axis), SDs (*y*-axis) and sample size (balanced) for the case of independent treatment and control group data (*ρ_CT_* = 0). Black dashed line indicates no effect (i.e., lnCVR = 0).

**Figure 4:**
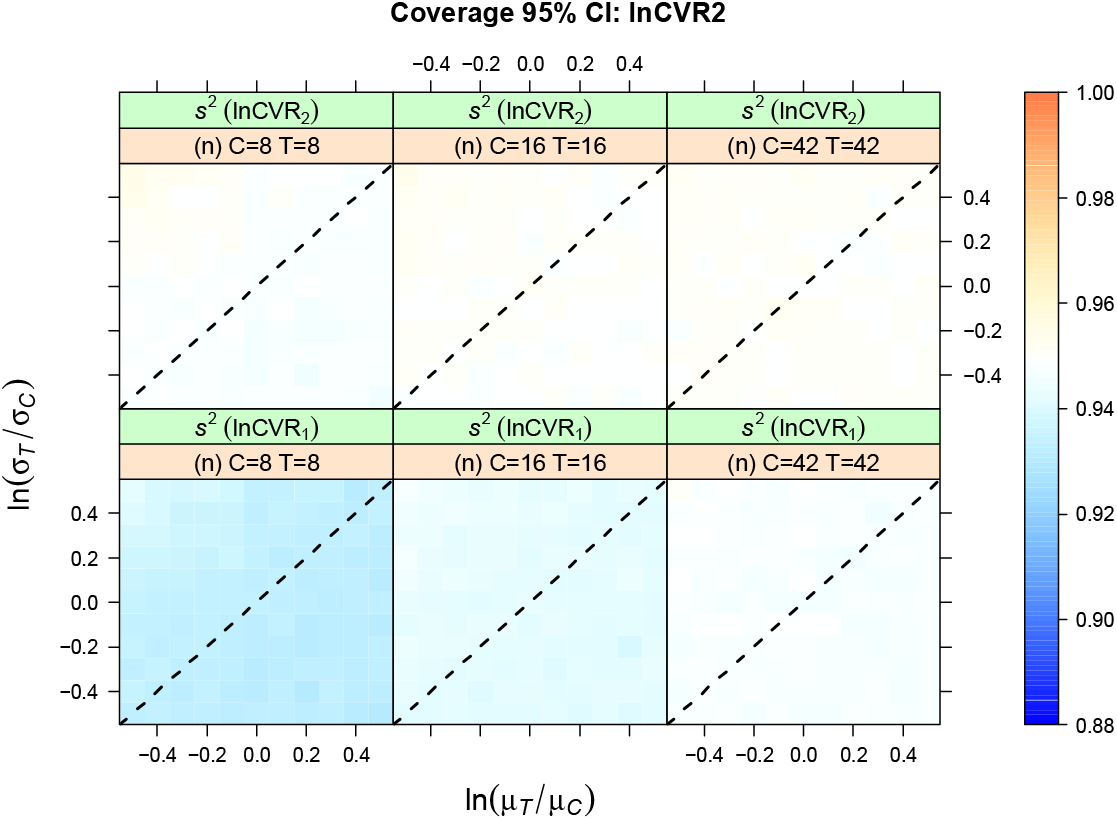
Coverage of 95% CIs based on estimators of the sampling variance of lnCVR as a function of the log ratio of population means (*x*-axis), SDs (*y*-axis) and sample size (balanced) for the case of independent treatment and control group data (*ρ_CT_* = 0). Black dashed line indicates no effect (i.e., lnCVR = 0).

For the case where treatment and control samples were dependent on one another (*ρ_CT_* = 0.8) lnCVR_4_ out-performed lnCVR_3_, with a pattern identical to that in Figure 1 (Figure S3). With regards the two estimators for dependent sampling variances, *s*^2^(lnCVR_3_) underestimated the variance where as *s*^2^(lnCVR_4_) overestimated the variance (Figure 5). These biases were within a reasonable range for larger samples, but were severe for small samples, and v(lnCVR_4_) in particular showed extreme upward bias (reaching 60% overestimate) when the SD of the treatment group differed from that of the control group (Figure 5). The CIs generated by v(lnCVR;) had a tendency to be too narrow whereas those generated by *s*^2^(lnCVR_4_) were too wide (Figure 6).

**Figure 5:**
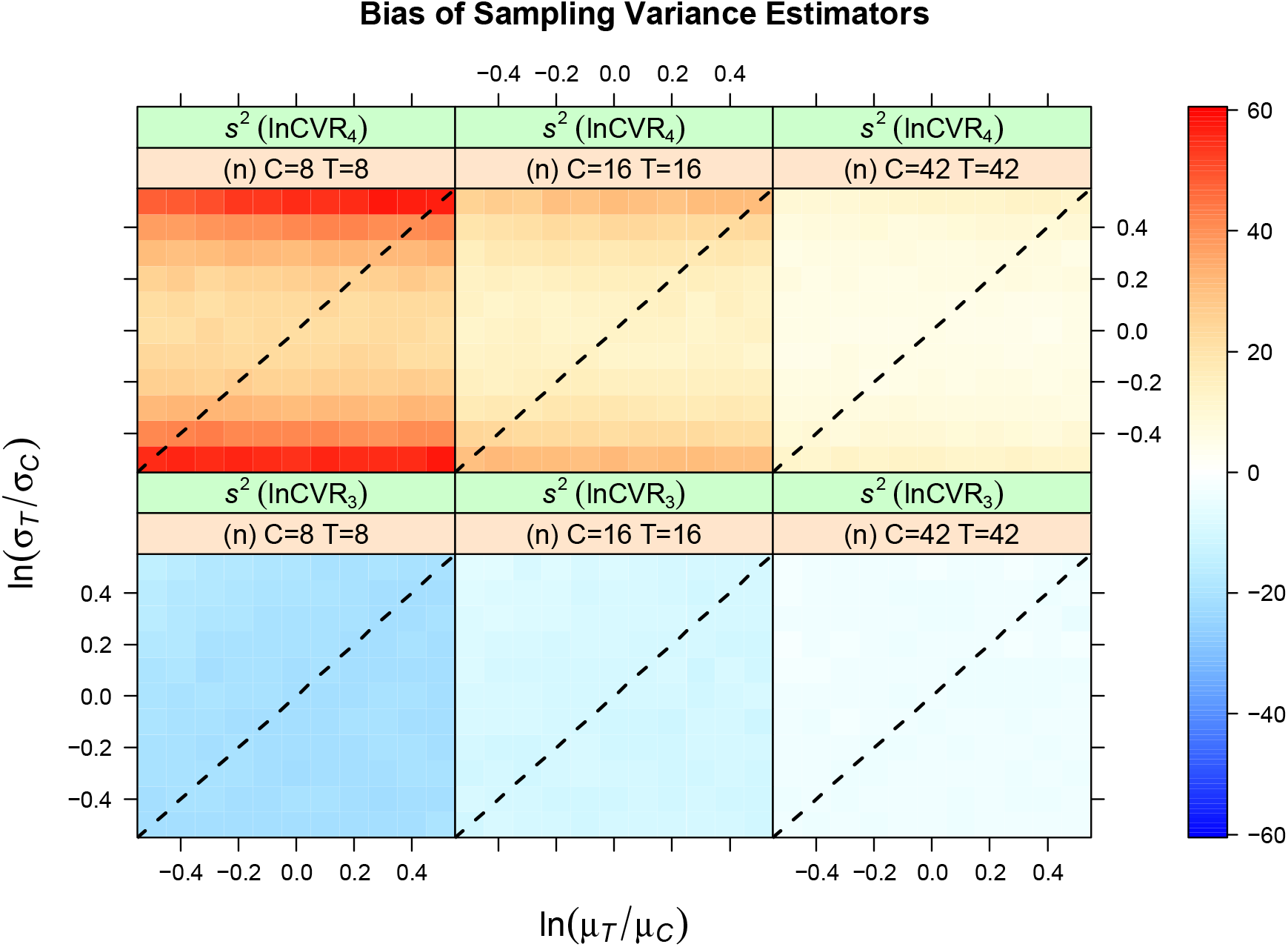
Bias in sampling variance estimators of lnCVR as a function of the log ratio of population means (*x*-axis), SDs (*y*-axis) and sample size (balanced) for the case of dependent treatment and control group data (*ρ_CT_* = 0.8). Black dashed line indicates no effect (i.e., lnCVR = 0).

**Figure 6:**
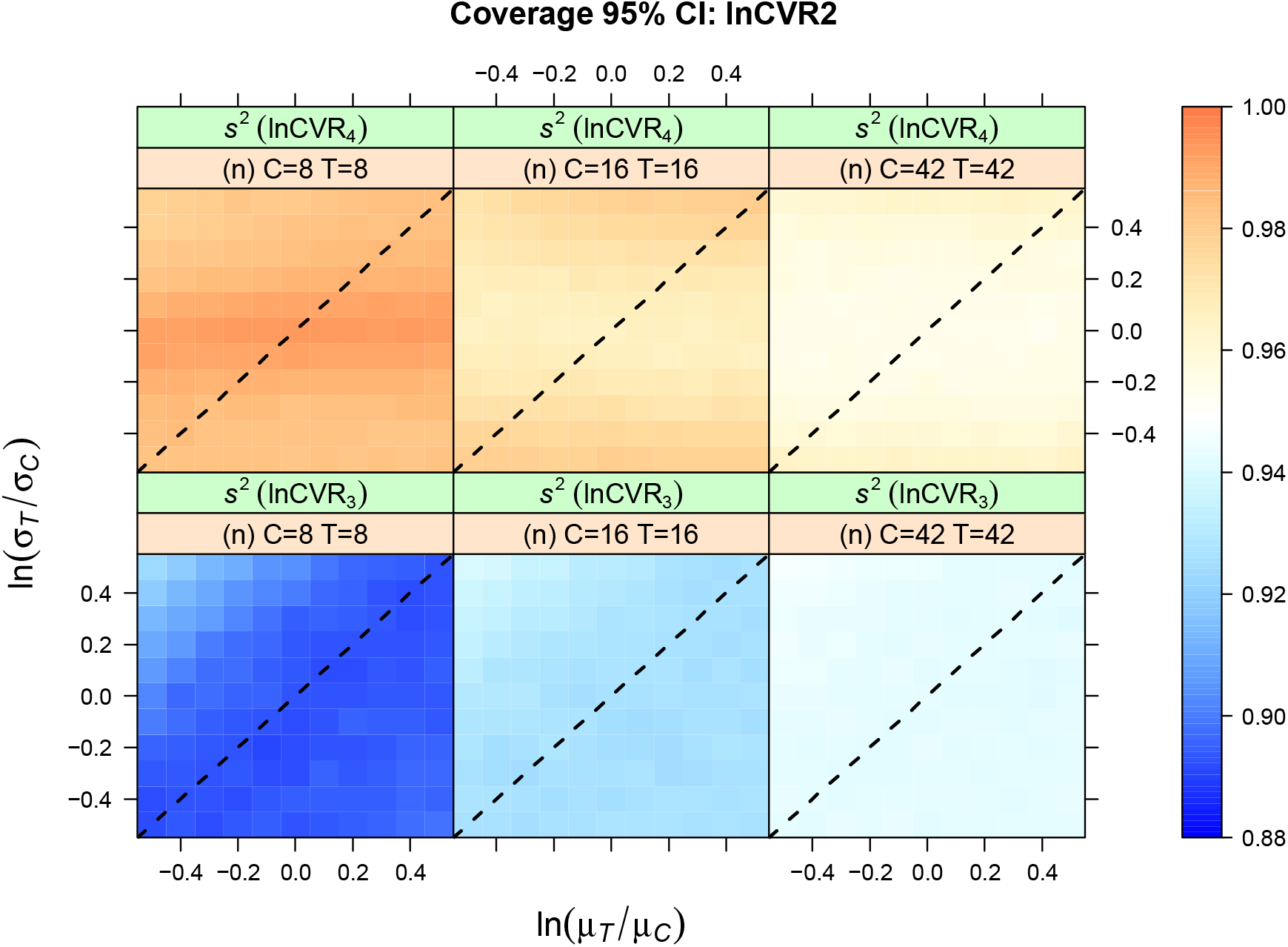
Coverage of 95% CIs based on estimators of the sampling variance of lnCVR as a function of the log ratio of population means (*x*-axis), SDs (*y*-axis) and sample size (balanced) for the case of dependent treatment and control group data (*p_CT_* = 0.8). Black dashed line indicates no effect (i.e., lnCVR = 0).

## 4. WORKED EXAMPLES

We now provide two examples: one from the field of ecology and the other from the health sciences. All meta-analytic models (random-effects meta-analysis) were fitted using the ‘rma’ function (with default settings) in *metafor*^32^

### 4.1 Example 1: Carbon dioxide levels and plant mass

Curtis, Wang^33^ performed a meta-analysis of experimental studies that tested for the effects of elevated carbon dioxide (CO_2_) levels on woody plant mass. Briefly, these studies compared the total biomass (above and below ground) of plants grown under ambient and artificially elevated (~100% increase) CO_2_ levels. Studies were performed in a range of contexts, including highly controlled (e.g., green houses) and less controlled (e.g., field sites) environments, as well as across temperature, light, water, and soil-fertility levels. Replication was at the level of the locale (e.g., plot/site/greenhouse) at which a treatment was applied, and treatment/control groups may be correlated (i.e., non-independent) if, for example, locales experiencing different treatments are paired spatially or temporally. However, the degree to which such correlations are present was not stated. Aggregating 102 effect sizes (lnRR), Curtis, Wang^33^ found that the mean biomass of woody plants at a site increases by, on average, 28.8% under elevated CO_2_ conditions. However, there was evidence that the effect is moderated by the presence of other stressors such as under nutrient- or light-limited conditions.

Here we ask whether elevated CO_2_ levels also increase among-replicate variability in plant biomass using lnCVR. We tested the sensitivity of the analysis to the assumption that treatment and control groups are uncorrelated. Because we do not know precisely which effect size data come from paired designs, we calculated effect sizes and sampling variance assuming complete independence (0% of effect sizes have correlated groups), varying degrees of partial dependence (a random subset of 20%, 60%, or 80% effect sizes have correlated groups; *r_CT_* = 0.8), or complete dependence (100% of effect sizes have correlated groups; *r_CT_* = 0.8). For those effect sizes that were assumed to be uncorrelated we used lnCVR_2_ and *s*^2^(lnCVR_2_), and for those that are correlated lnCVR_4_ and *s*^2^(lnCVR_3_).

There was evidence for a mean-variance relationship under both elevated and ambient CO_2_ levels (Figure 7A). The influence of increasing the percentage of effect sizes that are assumed to come from correlated groups on a random-effects meta-analysis is shown in Table 1. There are some qualitative differences in the interpretation of the overall effect, whereby the associated CI spans zero in some cases, but not others (Table 1). In all cases the sign of the overall effect is stable and suggests that elevating CO_2_ levels on average decreases the CV in biomass among replicates (possibly by somewhere between 100 × (1 − exp(−0.078) = 7.5 to 100 × (1 − exp(−0.116) = 10.9 percent). The effect of increasing the number of studies with correlated groups on the estimated inter-effect size heterogeneity, is however, much more dramatic. As less independence is assumed, the amount of heterogeneity (absolutely, in terms of 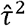, and relatively, in terms of *I*^2^) increases substantially (Table 1), such that when 40% or more of the studies are have used paired designs, Cochran’s *Q* test yields a significant result.

**Figure 7:**
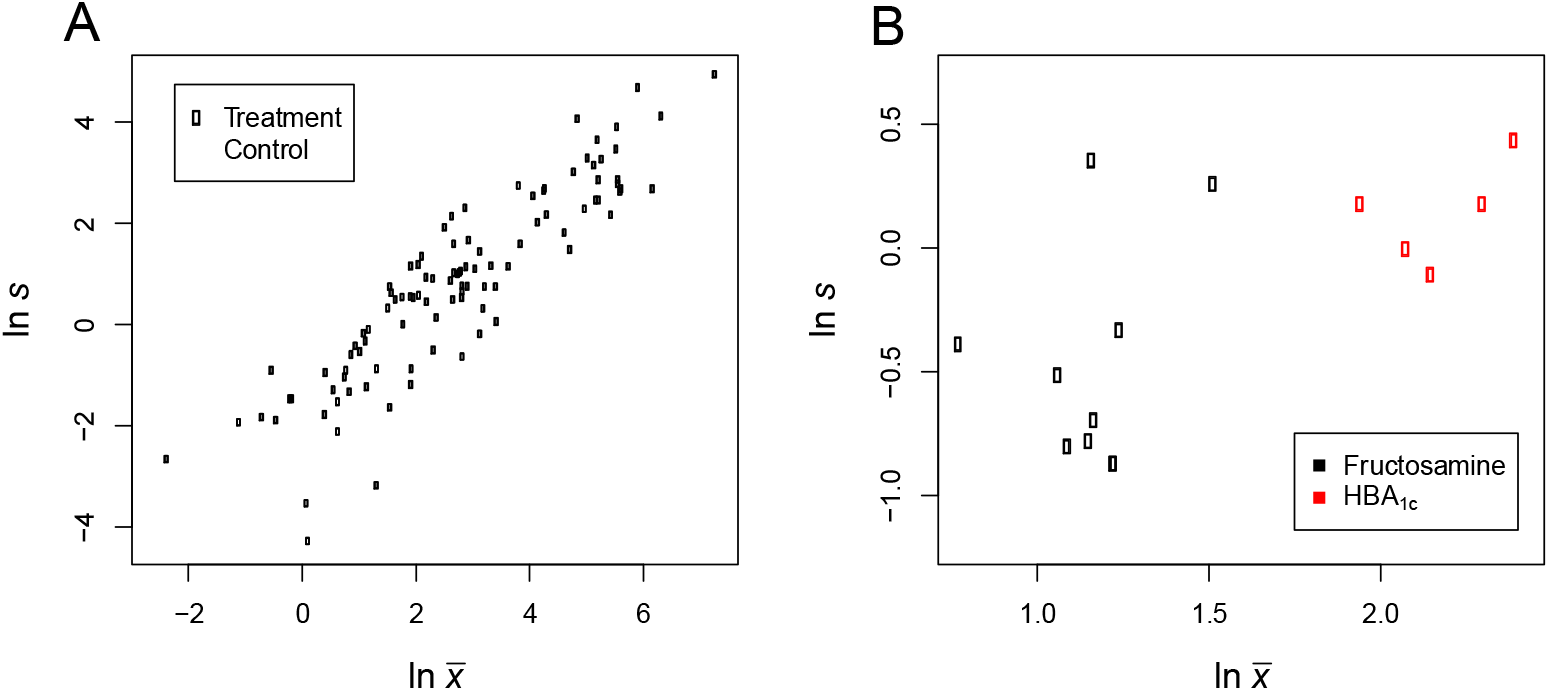
Association between log sample mean 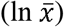 and log sample standard deviation (ln *s*) for treatment (hollow points) and control (solid points) groups in the data from; A) Curtis, Wang^33^, where the outcome is woody plant biomass under elevated (treatment) *vs* ambient (control) CO_2_ levels; and B) Brand-Miller et al. (2003) where the outcome is a measure of glycemia in diabetic individuals on low (treatment) *vs* high (control) glycemic index diets. Note in (B) measures of glycemia are either fructosamine (black points) or HbA1c (red points) levels, where lower levels indicate better gylcemic control.

**Table 1:** Estimates of the overall effect (lnCVR) and heterogeneity statistics from randomeffects meta-analyses of woody plant biomass under elevated *vs* ambient CO_2_ levels. Negative estimates indicate lower CV under elevated CO_2_. Models were refitted based on effect sizes assuming increasing % of the effect sizes contain correlated (dependent) treatment and control group data (*r_CT_* = 0.8). LCI and UCI indicate the lower and upper 95% confidence interval bounds. Data from Curtis, Wang^33^.

| % Correlated | Estimate | SE | LCI | UCI | $\tau^2$ | $I^2$ | $Q$ | $p(Q)$ |
| --- | --- | --- | --- | --- | --- | --- | --- | --- |
| 0 | -0.078 | 0.044 | -0.163 | 0.008 | 0.000 | 0.000 | 85.75 | 0.861 |
| 20 | -0.090 | 0.055 | -0.198 | 0.017 | 0.082 | 35.02 | 141.0 | 0.005 |
| 40 | -0.093 | 0.057 | -0.205 | 0.019 | 0.133 | 55.00 | 191.8 | <0.001 |
| 60 | -0.095 | 0.057 | -0.207 | 0.017 | 0.161 | 64.28 | 240.7 | <0.001 |
| 80 | -0.118 | 0.054 | -0.225 | -0.011 | 0.165 | 68.31 | 267.0 | <0.001 |
| 100 | -0.116 | 0.053 | -0.219 | -0.012 | 0.17 | 73.77 | 294.2 | <0.001 |

### 4.2 Example 2: Low glycemic index diets and glycemic control in diabetic subjects

Brand-Miller, Petocz, Hayne, Colagiuri^34^ performed a meta-analysis of studies designed to test the effects of low glycemic index (GI) diets on bio-markers of glycemic control in diabetic (type 1 and 2) individuals. Individuals were given either low or high GI diets, after which glycemia was measured using HbA_1c_ and/or fructosamine levels. These two markers quantify glycemia over longer *vs* shorter time periods respectively, where lower levels indicate better glycemic control. The studies differed somewhat in the overall GI of the diets used and the duration for which subjects were on the diets. The studies used a mixture of parallel designs where the individuals in each treatment group are completely independent, and cross-over designs where each individual was subject to both treatments. Brand-Miller, Petocz, Hayne, Colagiuri^34^ acknowledged that for those studies with a cross-over design, there will be a degree of correlation among the treatment and control condition data. They tested the sensitivity of their results to any such correlation by repeating the analyses assuming complete independence (*r_CT_* = 0) and also assuming that groups are correlated (*r_CT_* = 0.34; based on one of the studies in their primary literature). Their analyses of 14 effect sizes (mean differences, expressed in terms of percent; 11 from studies with cross-over designs) suggested that measures of glycemia are decreased by 6.8 percentage points (improved glycemic control) on low GI diets irrespective of their assumptions about correlations among groups. The authors used a fixed-effect meta-analytic model, and did not present heterogeneity statistics.

We tested whether low GI diets affect inter-individual variability in glycemic control using lnCVR. Unlike example 1, here we do know which studies contain dependent groups (those with cross-over designs), although the strength of the dependence is not precisely known. For independent designs we calculated effect sizes and sampling variances *via* lnCVR_2_, and *s*^2^(lnCVR_2_). For those studies using a cross-over design we calculated lnCVR_4_ and *s*^2^(lnCVR_3_) assuming treatment and control data are correlated with *r_CT_* = 0, 0.3, 0.5, and 0.8. Where more than one measure of glycemia was presented from a single study, we primarily use fructosamine levels (this being the more widely reported measure).

We observed a mean-variance relationship amongst both measures of glycemic control within the two treatment groups (Figure 7B). The results of random-effects meta-analyses fitted to the effect sizes are given in Table 2. The analyses estimated that on low-GI diets the CV in biomarkers of glycemic control is on average reduced by between 13% (100 × (1 − exp(−0.135)) and 18% (100 × (1 − exp(−0.177)) compared to high-GI diets. However, as the degree of correlation among data from cross-over trials increased, there was a marginal reduction in the overall effect magnitude and an increase in the associated SE (Table 2); for *r_CT_* = 0.5, the overall effect was not statistically significant. With increasing correlation, heterogeneity also increased (Table 2). Where we assumed complete independence (*r_CT_* = 0), there was no evidence for heterogeneity, but for *r_CT_* = 0.8, we detected inter effect size heterogeneity (Table 2).

**Table 2:** Estimates of overall effect (lnCVR) and heterogeneity from random-effects metaanalyses of glycemic control in diabetics on low- *vs* high-GI diets. Negative estimates indicate lower CV on a low-GI diet. Models were refitted from effect sizes assuming differing strength of correlation (*r_CT_*) among repeated measured from the same individuals in cross-over trials. LCI and UCI indicate the lower and upper 95% confidence interval bounds. Data from Brand-Miller, Petocz, Hayne, Colagiuri^34^.

| $r_{CT}$ | Estimate | SE | LCI | UCI | $\tau^2$ | $I^2$ | $Q$ | $p(Q)$ |
| --- | --- | --- | --- | --- | --- | --- | --- | --- |
| 0 | -0.177 | 0.070 | -0.314 | -0.039 | <0.001 | 0.006 | 15.88 | 0.321 |
| 0.3 | -0.162 | 0.075 | -0.308 | -0.015 | 0.012 | 15.14 | 18.92 | 0.168 |
| 0.5 | -0.151 | 0.080 | -0.307 | 0.006 | 0.030 | 32.73 | 22.33 | 0.072 |
| 0.8 | -0.135 | 0.091 | -0.314 | 0.044 | 0.085 | 70.44 | 42.58 | <0.001 |

## 5. DISCUSSION AND CONCLUSIONS

We recommend that meta-analysts use the following estimator of the lnCVR for independent study designs:

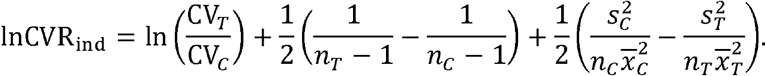

For dependent study designs we recommend the use of the following point estimator:

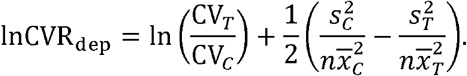

Under the simulated conditions explored, these estimators exhibited minimal bias, where ‘naïve’ estimators displayed systematic biases, substantially overestimating large positive effects, especially when sample sizes were small. Compared to previous estimators^15^, this revision contains an additional term, 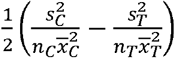, which has also been shown to reduce bias in mean effects estimated *via* lnRR^23^. We also recommend that the following estimators for the sampling variance of lnCVR be used for independent and dependent study designs, respectively:

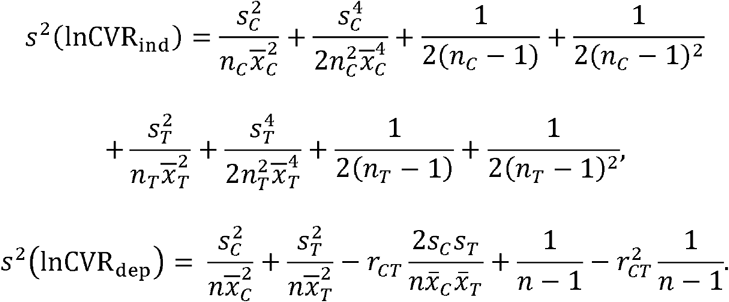

Our simulations demonstrate that the estimator for independent designs performs very well and 95% CIs based on a *z* distribution give coverage at the nominal level. The estimator for dependent cases slightly underestimates the actual sampling variance in lnCVR, and will generate CIs (based on *z* or *t* distributions) that are slightly too narrow. This might be due to the substitution of *r_CT_* for the unknown true correlation in the equation for the sampling variance without further account of the additional source of uncertainty this introduces. CIs that are too narrow may be more troublesome in that they can lead to inflated type-1 error rates (a more conservative estimator, *s*^2^(lnCVR_4_), is given above, although this approach may substantially overestimate the sampling variance for small samples). Note that these recommended estimators are now available in the ‘escalc’ function in the development version of *metafor* (https://github.com/wviechtb/metafor), and will eventually be implemented in the CRAN version.

We used the recommended estimators to evaluate whether: 1) increased CO_2_ levels affect variation in woody plant biomass, and 2) low-GI diets alter between-individual variation in glycemic control in diabetics. In both cases, we found that the treatments have a tendency to decrease the CV. In both cases the analyses were sensitive to assumptions about the degree to which treatment and control data are correlated. Assuming higher degrees of correlation resulted in small changes in the overall effect (and its standard error). Although these parameters were relatively stable, for estimates with CIs close to zero, changing assumptions about group independence can affect inference. Increasing the degree of correlation dramatically increased the estimated between-effect size heterogeneity, which could change conclusions about the consistency of the reported effects. This trend can be explained by the fact that as more/stronger correlations are assumed the sampling variances associated with the individual effect sizes shrink, effects are assumed to be more precise, and sampling variability therefore becomes less able to explain the variation among the effects. Our results corroborate the points made by Becker^28^, who introduced an estimator for the sampling variance of SMD for dependent groups.

As is the case with any exercise in data analysis, the most appropriate technique to use will depend on the question being asked. Where the analyst is able to determine with a reasonable degree of certainty that a mean-variance relationship does not exist, lnVR may be a more useful measure of between-group differences in variability than lnCVR. This is because lnCVR risks conflating effects on the SD with effects on the mean. In other instances, the user may be more interested in ascertaining whether a treatment alters the SD irrespective of a mean-variance relationship (e.g., in questions related to power and study design), and again lnVR would be an appropriate choice. However, where mean-variance relationships are present, and the analyst is interested in whether the variation is greater/lower than expected given the mean, lnCVR is useful. For some matters, it may even be common practice for the primary literature to describe variation in terms of CV rather than SD. For instance, in ecology and evolution it is common to present CV when comparing variability amongst species/traits that exist on different scales because CV is a relative measure^35^. We note that such a practice is not necessarily required for meta-analysis because lnVR is also a relative measure of variation, and as such should also do a good job of correcting for inter-system differences in scale. Nevertheless, where CV is the measure of variability commonly reported in the primary literature, the user may find it intuitive (or even necessary) to use lnCVR.

Nakagawa, Poulin, Mengersen, et al.^15^ also present alternative arm-based models (and discuss bivariate models) for meta-analysis of variation. The lnCVR metric assumes that changes in the mean are associated with proportional changes in the SD. Arm-based (and bivariate) models are an alternative for meta-analysis which allow the user to circumvent the assumption of proportionality. Arm-based models, however, are not without their critics who argue that these methods are radical departure from established meta-analytic thinking (see^16^). Like other (contrast-based) effect size measures that reflect the difference between two groups (e.g., the standardized mean difference, log response ratio, log risk/odds ratio or the risk difference), lnCVR readily integrates with our most widespread analytical paradigms, offering a convenient and intuitive method for meta-analysis of variability.

Finally, we finish by reiterating the point made by Nakagawa, Poulin, Mengersen, et al.^15^, and echoed by subsequent papers using lnCVR in different fields of study^17–21^. Meta-analysis of variation can tackle entirely new questions and open our eyes to insights that are hidden in datasets. The datasets required to gain these insights already exist because lnCVR is based on the same summary statistics as SMD and lnRR; means, SDs, and sample sizes. We suspect over 50,000 datasets of this sort have already been collected (c.f.^36^). In this regard it is vital that meta-analytic ‘raw’ data are made available and reusable in the spirit of open and transparent science^37,38^.

## ACKNOWLEDGEMENTS

AMS was supported by an Australian Research Council Discovery Early Career Researcher Award (ARC DECRA: DE180101520). SN was supported by an ARC Discovery Grant (DP180100818). We would like to thank Professor Jennie Brand-Miller for her thoughts on our analysis of glycemia in diabetic individuals.

