## Supplementary Materials S1 for "Revisiting and expanding the meta-analysis of variation: The log coefficient of variation ratio, lnCVR"

**Text S1**

It has been proposed that an unbiased estimator for the population CV is $\left( 1+\frac{1}{4n} \right)\mathrm{CV}$ ^1^. Based on this, lnCVR could be estimated as:

$$\ln\mathrm{CVR}=\ln\left( \left[ 1+\frac{1}{4n_{T}} \right]\mathrm{CV}_{T} \right)-\ln\left( \left[ 1+\frac{1}{4n_{C}} \right]\mathrm{CV}_{C} \right).$$

Note, however, that this is not automatically an unbiased estimator. Even if $\left( 1+\frac{1}{4n} \right)\mathrm{CV}$ was unbiased for the true CV (which is not the case, as this is just a rough approximation), this does not imply that $\left( \left( 1+\frac{1}{4n_{T}} \right)CV_{T} \right)/\left( \left( 1+\frac{1}{4n_{C}} \right)CV_{C} \right)$ is unbiased (and conversely, even though CV is biased, this does not automatically imply that $CV_{T}/CV_{C}$ or $\mathrm{lnCV}R_{1}$ as given in the main text are biased). Additionally, it is unclear what the appropriate sampling variance for the above estimator should be. An estimator of sampling variance for $\left( 1+\frac{1}{4n} \right)\mathrm{CV}$ is given in ^1^, but is unsupported by any derivation or citation. An alternative publication gives some derivation for the required sampling variance ^2^, but this can be considered at best a rough approximation and performs poorly under simulation.

**Text S2**

Based on the ‘second-order’ sampling variance estimators, when two (or more) treatment groups share a common control group (i.e., shared control) ^3^, it is possible to estimate the sampling covariance. When we have treatment A and treatment B, the sampling covariance between lnRR^A^ and lnRR^B^ is (c.f. ^4^ REF):

$$COV\left( \mathrm{lnRR}^{A}, \mathrm{lnRR}^{B} \right)=\frac{s_{C}^{2}}{n_{C}\overline{x}_{C}^{2}}+\frac{s_{C}^{4}}{2n_{C}^{2}\overline{x}_{C}^{4}}.$$

Similarly, the sampling covariance between lnVR^A^ and lnVR^B^ and the one between lnCVR^A^ and lnCVR^B^ are respectively:

$$COV\left( \mathrm{lnVR}^{A}, \mathrm{lnVR}^{B} \right)=\frac{1}{2}\left( \frac{1}{n_{C}-1}+\frac{1}{\left( n_{C}-1 \right)^{2}} \right),$$

$$COV\left( \mathrm{lnCVR}^{A}, \mathrm{lnCVR}^{B} \right)=\frac{s_{C}^{2}}{n_{C}\overline{x}_{C}^{2}}+\frac{s_{C}^{4}}{2n_{C}^{2}\overline{x}_{C}^{4}}+\frac{1}{2\left( n_{C}-1 \right)}+\frac{1}{2\left( n_{C}-1 \right)^{2}}.$$

These covariance estimators can be used by adding a variance-covariance matrix in meta-analytic ^3^

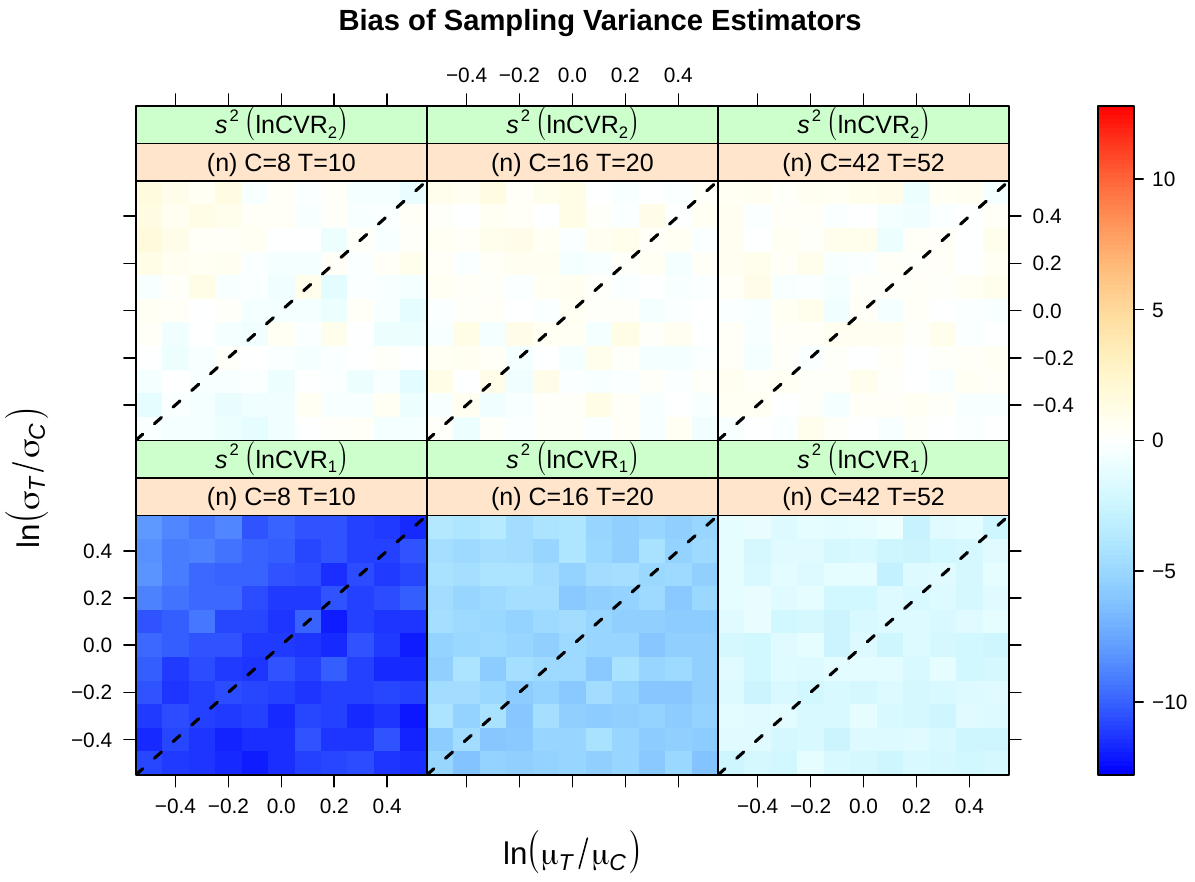

**Figure S1:** Bias in sampling variance estimators of lnCVR as a function of the log ratio of population means (*x*-axis), SDs (*y*-axis), and sample size (unbalanced) for the case of independent treatment and control group data ($\rho_{CT}$ = 0). Black dashed line indicates no effect (i.e., lnCVR = 0).

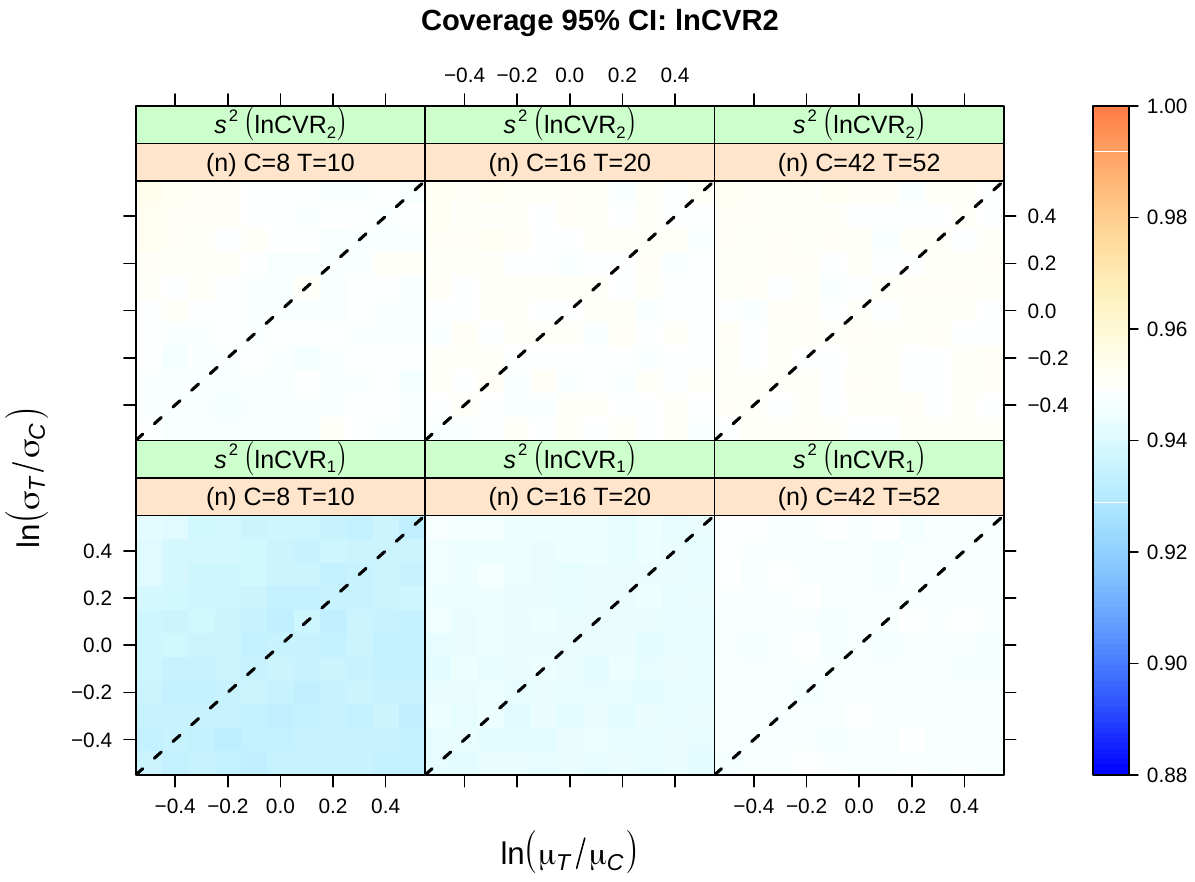

**Figure S2:** Coverage of 95% CIs based on estimators of the sampling variance of lnCVR as a function of the log ratio of population means (*x*-axis), SDs (*y*-axis), and sample size (unbalanced) for the case of independent treatment and control group data ($\rho_{CT}$ = 0). Black dashed line indicates no effect (i.e., lnCVR = 0).

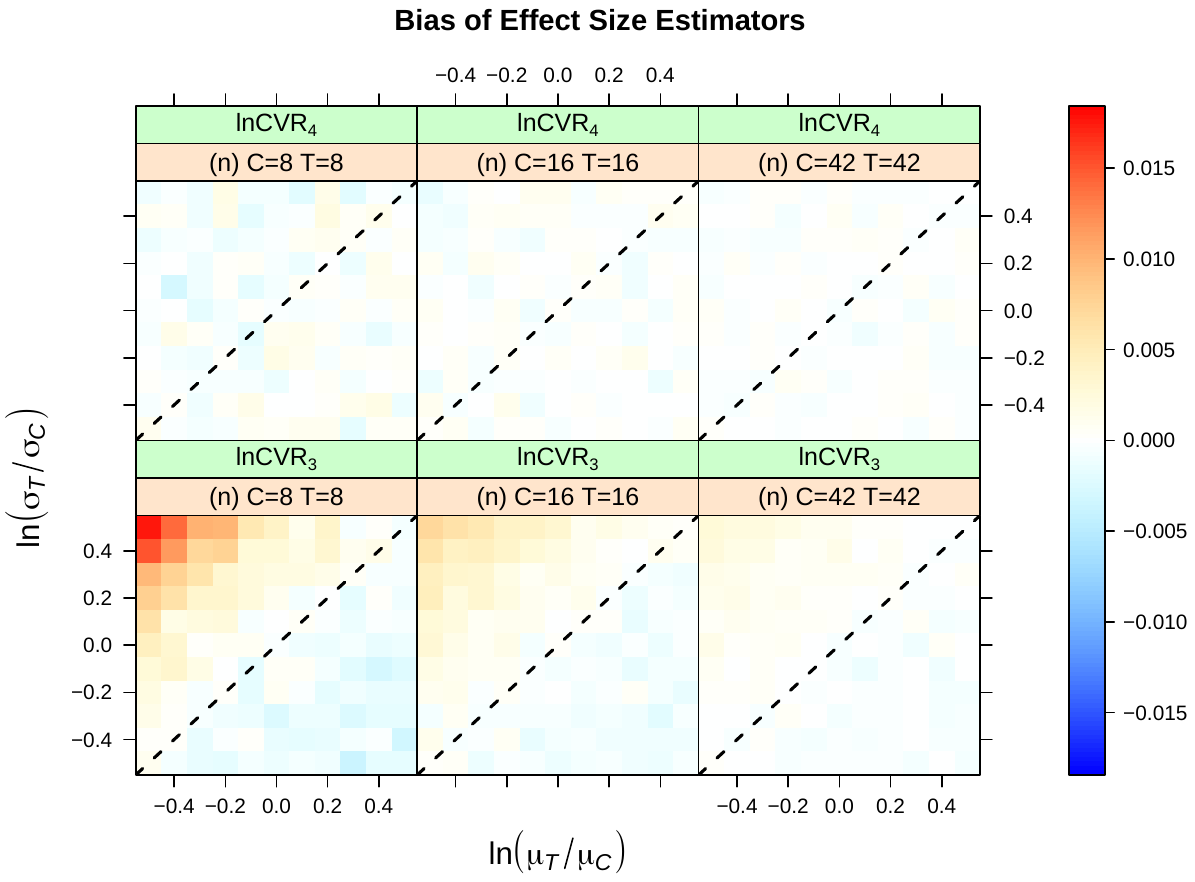

**Figure S3:** Bias in effect size estimators of lnCVR as a function of the log ratio of population means (*x*-axis), SDs (*y*-axis), and sample size (balanced) for the case of dependent treatment and control group data ($\rho_{CT}$ = 0.8). Black dashed line indicates no effect (i.e., lnCVR = 0).
